# Rodent ultrasonic vocal interaction resolved with millimeter precision using hybrid beamforming

**DOI:** 10.1101/2023.01.18.524540

**Authors:** M. L. Sterling, B. Englitz

## Abstract

Ultrasonic vocalizations (USVs) fulfill an important role in communication and navigation in many species. Because of their social and affective significance, rodent USVs are increasingly used as a behavioral measure in neurodevelopmental and neurolinguistic research. Reliably attributing USVs to their emitter during close interactions has emerged as a difficult, key challenge. If addressed, all subsequent analyses gain substantial confidence.

We present a hybrid ultrasonic tracking system, HyVL, that synergistically integrates a high-resolution acoustic camera with high-quality ultrasonic microphones. HyVL is the first to achieve millimeter precision (~3.4-4.8mm, 91% assigned) in localizing USVs, ~3x better than other systems, approaching the physical limits (mouse snout ~ 10mm).

We analyze mouse courtship interactions and demonstrate that males and females vocalize in starkly different relative spatial positions, and that the fraction of female vocalizations has likely been overestimated previously due to imprecise localization. Further, we find that male mice vocalize more intensely when interacting with two mice, an effect mostly driven by the dominant male.

HyVL substantially improves the precision with which social communication between rodents can be studied. It is also affordable, open-source, easy to set up, can be integrated with existing setups, and reduces the required number of experiments and animals.

**Data & Code Availability:** During the review process, reviewers can access all Data and Code via the link below: https://data.donders.ru.nl/loqin/reviewer-208072048/iJ4c-oRNlPIp3vArKiYQ0lAW9FipiHL8foxSzwt1FDAUpon acceptance, these materials will be made available to the public.

## Introduction

Ultrasonic vocalizations (USVs) fulfill an important role in animal ecology as means of communication or navigation in many rodents^1–5^, bats^6^, frogs^7^, cetaceans^8^, and even some primates.^9,10^ In many of these species, USVs have been shown to be present innately and to have significance at multiple stages of life, from neonates^11^ to adults^1^, often with diverse functions as distress/alarm calls^11,12^, courtship signals^13^, territorial defense signals^14^, private communication^10^, and echolocation^6^. USVs have been extensively studied in mice, where their communicative significance has been widely demonstrated by their influence on conspecific behavior^15–20^ (also in line with observational studies^21–24^). USVs can be grouped into different types that are highly context-dependent^17,18,22,25–37^, and USV syntax itself is predictive of USV sequence.^38^ Taken together, the current literature suggests USVs convey affective and social information in different behavioral contexts. This is further supported by the modulatory effect that testosterone and oxytocin have on USV production.^39–45^ Importantly, the neuronal circuitry underlying USVs has recently been identified and is being studied extensively.^20,25,46–53^

Because of their social and affective significance and our growing mechanistic understanding, mouse USVs are increasingly being used as a behavioral measure in neurodevelopmental and neurolinguistic translational research.^26,32,49,54–59^ Their manipulation and precise measurement not only provide the basis for tackling many fundamental questions but also pave the way, via advanced animal models, for the discovery of essential, novel drug targets for many debilitating conditions such as autism-spectrum disorder^58,60^, Parkinson’s disease^61^, stroke-induced aphasia^62^, epilepsy aphasia syndromes^63^, progressive language disorders^64^, chronic pain^65^, and depression/anxiety disorders^66^, where ultrasonic vocalizations serve as a biomarker for animal well-being and normal development. Consequently, we expect the scientific importance of mouse USVs to continue to increase in the coming years, highlighting the necessity to advance the methods required for their study. In recent years, substantial advances have been made in USV detection^67–71^, classification^67,68,70,72^, and localization.^73–76^

Localization is of particular importance during social interactions, when most USVs are emitted and any meaningful analysis of USV properties rests on a reliable assignment of each USV to its emitter. This task is complex for multiple reasons: (i) most USVs are emitted at close range, (ii) social behavior often requires free movement of the animals, and (iii) USV production is invisible.^37,77^ With reliable assignment, all subsequent analyses can be conducted with substantial confidence concerning each USV’s emitter. Although USVs could in theory be classified and assigned based on their shape^13,78–81^, this approach will depend strongly on different behavioral contexts and strains. Recent advances in acoustic localization^74–76^ have improved the localization accuracy to 11-14 mm, however, close-up snout-snout interactions - which is when a large fraction of USVs are emitted - require an even higher precision.

We have developed an advanced localization system for USVs in which a high resolution ‘acoustic camera’ consisting of 64 ultrasound microphones with an array of 4 high-quality ultrasound microphones. Both systems can individually localize USVs but exhibit rather complementary patterns of localization errors. We fuse them into a hybrid system that exploits their respective advantages in sensitivity, detection, and localization accuracy. We achieve a median absolute localization error of 3.4-4.8 mm, translating to an assignment rate of ~91%. Compared to the previous state of the art^73,75^, the accuracy represents a three-fold improvement that halves the proportion of previously unassigned USVs. Given the physical dimensions of the mouse snout (ø~10 mm), this likely approaches the physical limit of localizability for USVs. We successfully apply it to and analyze dyadic and triadic courtship interactions between male and female mice. We demonstrate that the fraction of female vocalizations has likely been overestimated in previous analyses, due to a lack of precision in sound localization.

## Results

We analyzed courtship interactions of mice in dyadic and triadic pairings. The mice interacted on an elevated platform inside an anechoic booth (see *Fig*. 1A, for details see *Materials & Methods: Recording Setup*). Each trial consisted of 8 minutes of free interaction while movements were tracked with a high-speed camera (see *Fig*. 1B), and ultrasonic vocalizations (USVs) were recorded with a hybrid acoustic system composed of 4 high-quality microphones (i.e., USM4) as well as a 64-channel microphone array *(Cam64*, often referred to as an acoustic camera; see *Fig*. 1C for raw data samples, green and red dots mark the start and stop times of USVs).

**Figure 1:**
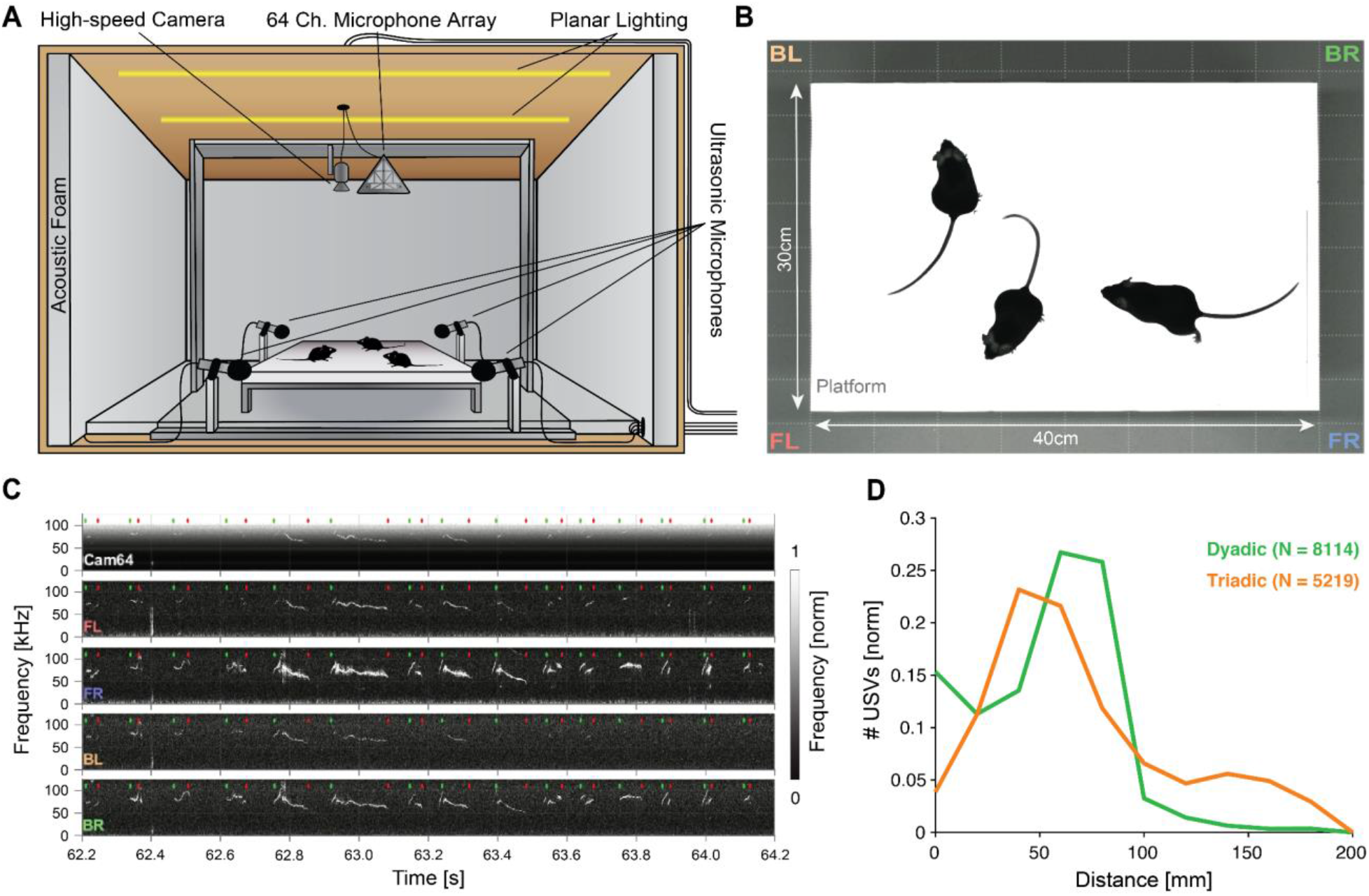
Mice emit ultrasonic vocalizations (USVs) in close proximity during courtship behavior. **A** Two or three mice of different sexes were allowed to interact freely on an elevated platform. Vocalizations were recorded with 4 high-quality ultrasonic microphones in a rectangular arrangement around the platform and a 64-channel microphone array (‘Cam64’) mounted above the platform. The spatial location of the pair was recorded visually with a high-speed camera. The platform was located in an ultrasonically sound-proof and anechoic box and illuminated uniformly using an array of LEDs. **B** Sample image from the camera that shows the high contrast between the mice and the interaction platform. The two-letter abbreviations indicate the locations of the 4 high-quality microphones (F = front, B = back, L = left, R = right). **C** Sample spectrograms from the four ultrasonic microphones and the average of all Cam64 microphones for a bout of vocalizations (start/end times marked by green/red dots). The Cam64 microphones are of lower quality than the USM4 microphones, evidenced by the rising noise floor for higher frequencies, affecting very high frequency USVs. **D** Most USVs in the present paradigm were emitted in close proximity to the interaction partners, with the vast majority within 10 cm snout-snout distance (i.e., ~93 and 72% for dyadic and triadic, respectively).

Most USVs were emitted in close proximity in dyadic and triadic pairings (see *Fig*. 1D). Reliably assigning most USVs to their emitter therefore requires a highly precise acousto-optical localization system. The presently developed Hybrid Vocalization Localizer (HyVL) system is the first to achieve sub-centimeter precision, i.e. ~3.4-4.8 mm (see *Fig*. 2 for an overview). This accuracy on the acoustic side is achieved by combining the complementary strengths of the USM4 and Cam64 data. The Cam64 data is processed using acoustic beamforming^82^ which delivers highly precise estimates (MAE = ~4-5 mm), but is not sensitive enough for very high-frequency USVs (see Fig. S2). The USM4 data is analyzed using our SLIM algorithm,^73^ which delivers accurate (MAE = ~11-14 mm) and less frequency-limited estimates. The methods exhibit a complementary pattern of localization errors, which predestines them for high synergy when combined (see below).

**Figure 2:**
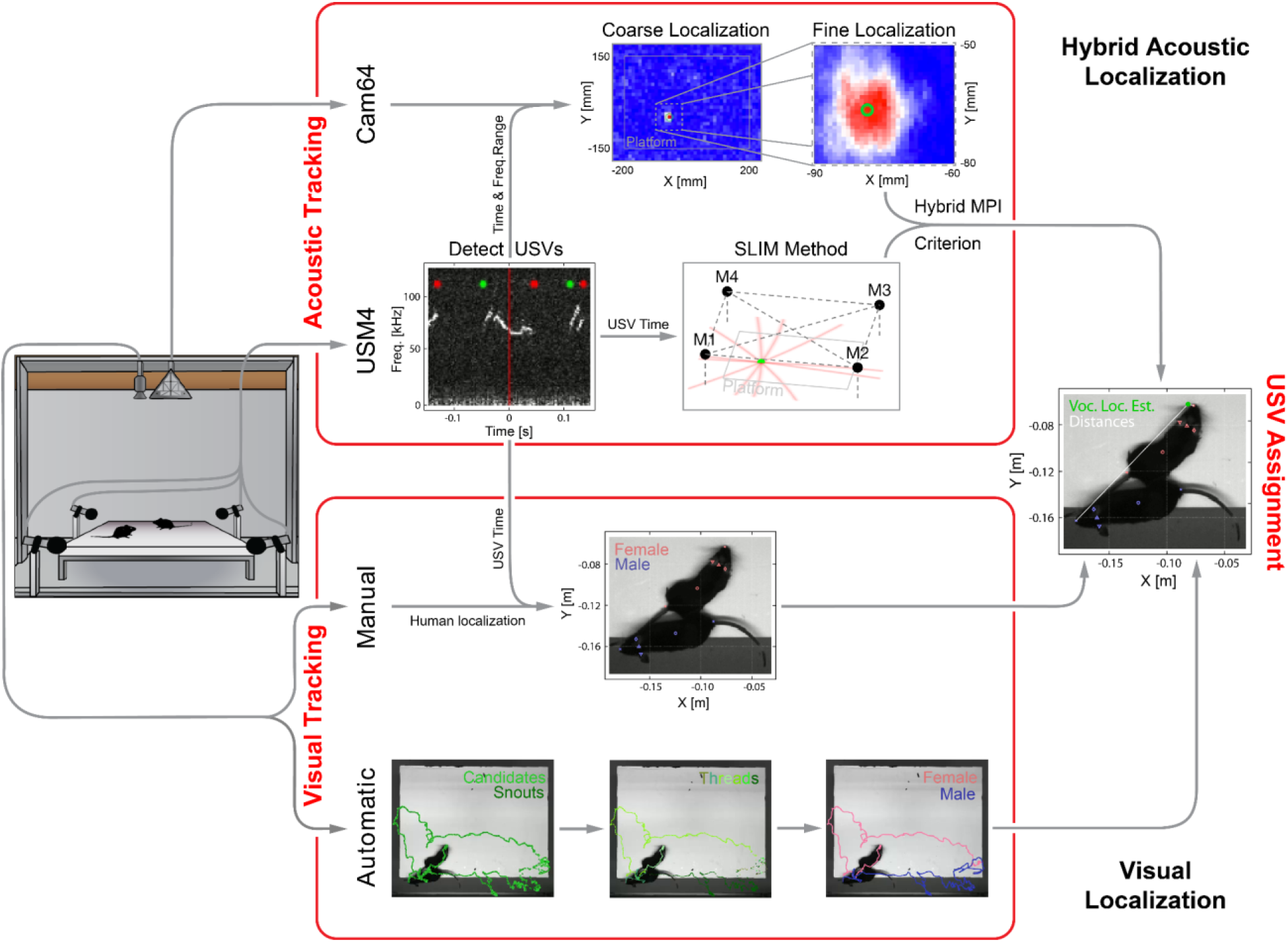
Overview of the combined acoustic and visual tracking pipeline. (Top) Acoustic tracking of animal vocalizations was enabled by a hybrid acoustic system, which recorded the sounds in the booth using a 64-channel ultrasonic microphone array (‘Cam64’) and 4 high-quality ultrasonic microphones (‘USM4’). Vocalizations were automatically detected using USM4 data (start/end times marked by green/red dots) and then localized on the platform using both the SLIM algorithm on USM4 data and delay-and-sum beamforming on the corresponding Cam64 data. The Cam64 localization proceeded in two steps: first coarse (10 mm resolution), then fine centered around the coarse peak at 1 mm resolution (30 x 30 mm local window). The local, weighted average (green circle) was then used as the USV origin localized by Cam64. For each USV, the Cam64 localization was chosen if its SNR > 5, otherwise the USM4/SLIM estimate was used (for details, see *Materials & Methods: Localization of Ultrasonic Vocalizations)*. (Bottom) Animals were tracked visually on the basis of concurrently acquired videos (50 FPS). Two tracking strategies were employed: (i) manual tracking in the video frames corresponding to the mid-point of USVs in all recordings and (ii) automatic tracking for all frames in dyadic recordings. (i) *Manual visual tracking:* the observer was presented with a combined display of the vocalization spectrogram and the concurrent video image at the temporal midpoint of each USV and annotated the snout and head center (i.e., midpoint between the ears). (ii) *Automatic visual tracking:* Started with finding the optimal locations of each marker based on marker estimate clouds produced by *DeepLabCut* (DLC; see Mathis et al., 2018) for all frames. Next, these marker positions were assembled into spatiotemporal threads with the same, unknown identity based on a combination of spatial and temporal analysis. Finally, the thread ends still loose were connected based on quadratic spatial trajectory estimates for each marker, yielding the complete track for both mice (see *Materials & Methods: Automatic Visual Animal Tracking* and *Suppl. Fig*. 1).

For each USV, a choice is made between the USM4/SLIM and Cam64/Beamforming estimates based on a comparison of each method’s USV-specific certainty and the relative position of the mice to the estimates, using an extended, hybrid Mouse Probability Index (MPI^76^). HyVL is the first system of its kind that exploits a hybrid microphone array to overcome the limitations of each subarray. The positions of the mice are obtained via manual and automatic video tracking using *DeepLabCut*,^83^ each of which achieve millimeter precision for localizing the snout.

Overall, 83 recordings were collected from 14 male and 4 female mice. Of these recordings, 55 were dyadic and 28 triadic. In all trials combined, 13406 USVs were detected.

### Precision of USV Localization

Assigning USVs to individual mice required combining high-speed video imaging with the HyVL location estimates at the times of vocalization. We manually tracked the animal snouts at the temporal midpoint of each USV to obtain near-optimal ground truth position estimates (see *Fig*. 2). We first assessed the relative structure of the localization errors between both methods, USM4/SLIM (*Fig*. 3A, green) and Cam64/Beamforming (*red*, each dot is a USV). While most errors were small, and clustered close to the origin of the graph (evidenced by the small Median Absolute Errors (MAE), shown as horizontal and vertical lines, respectively), the less frequent, larger errors exhibited an L-shape. This error pattern is an optimal situation for combining estimates from the two methods, to compensate for each other’s limitations. While the Cam64 data can compensate for single microphone noise through the large number of microphones, the nature of its micro-electromechanical systems (MEMS) microphones deteriorates for very high frequencies (see *Fig*. S2B). Conversely, the USM4 microphones show an excellent noise level across frequencies (see Fig. S2A) but can produce erroneous estimates if there is noise in a single microphone and have an intrinsic limitation in spatial accuracy due to the physical size of their receptive membrane (ø~20 mm).

**Figure 3:**
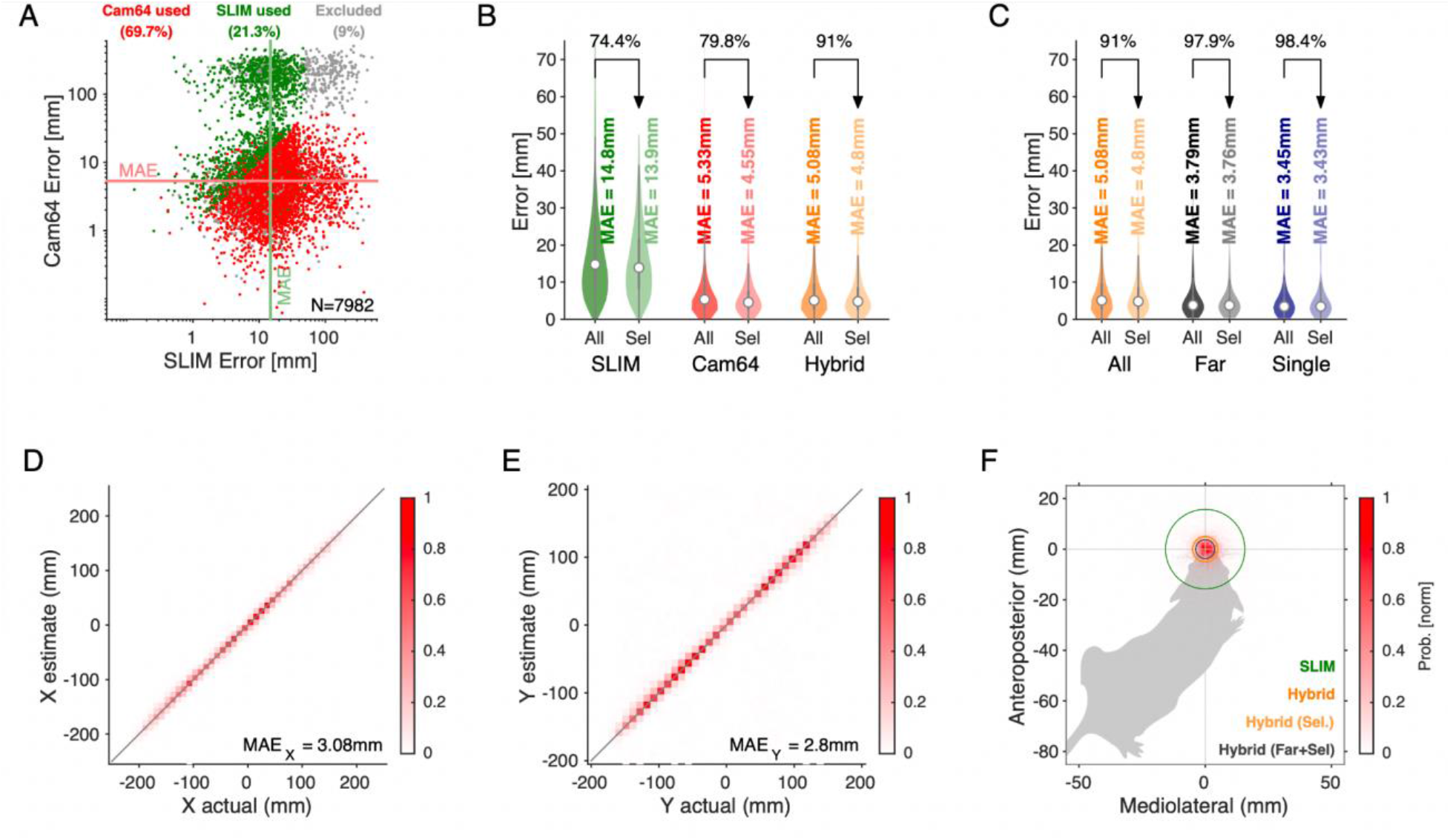
Spatial accuracy of localizing USVs during mouse social interaction improves ~3-fold over the state of the art.^73^. **A** The vast majority of USVs is localized with very small errors for both methods, concentrated close to the axes and thus hardly visible, evidenced by the median average errors (MAE) for Cam64 (light red line) and SLIM (light green line). The fewer larger errors form an L-shape, emphasizing the synergy of a hybrid approach that compensates for the weaknesses of each method. Location estimates were excluded (gray) if they were >50mm from either mouse, or the hybrid MPI<0.95. **B** The hybrid localization system HyVL (orange) combines the virtues of SLIM and Cam64 enabling the localization of 91.1% of all USVs (light orange), achieving an MAE = 4.8 mm. Cam64 localization (red) alone only includes 74.4% of all USVs, but at an MAE = 4.5 mm (light red). SLIM-based localization (green) only includes 80.0% of all USVs, at an MAE = 14.6 mm (light green, see *Materials & Methods: USV Assignment* for details on the relation between accuracy and selection criteria). **C** USVs emitted when all animals were >100 mm apart and a single mouse condition was used to assess the ideal accuracy of HyVL. For the far condition, virtually all USVs (332/339, 97.9%) were assigned at an MAE = 3.8 mm, similarly to the single animal condition (MAE = 3.4 mm, 251/255, 98.4%). **D/E** Comparison of actual with estimated snout locations along the X (horizontal; **D**) and Y (vertical; **E**) dimensions indicating strong agreement. Colors indicate peak-normalized occurrence rates. **F** Centered overlay of USV localizations relative to emitter snout. Precision is depicted as a circle with a radius equivalent to the median absolute error (green: SLIM; orange: HyVL, all USVs; light orange: HyVL, selected USVs, dark gray: HyVL, when mice >100 mm apart).

We therefore designed an analytical strategy to combine the estimates of both systems to optimize the number of reliably assignable USVs, while evaluating the resulting spatial accuracy alongside. Briefly, the location estimates of both methods each come with an estimate of localization uncertainty. First, we assess for each method’s estimate how reliably it can be assigned to one of the mice, taking into account the positions of the other mice. This is quantified using the Mouse Probability Index (MPI),^76^ which compares the probability of assignment to a particular mouse to the sum of probabilities for all mice, weighted by the estimate’s uncertainty. If the largest MPI exceeds 0.95, it is considered a reliable assignment to the corresponding mouse. If both methods allowed reliable assignments, the one with smaller residual distance was chosen. If only one method was reliable for a particular USV, its estimate was used. If neither method allowed for reliable assignment, the USV was not used for further analysis. This typically happens if the snouts are extremely close or the USV is very quiet. This approach outperformed many other combination approaches in accuracy and assignment percentage, e.g. Maximum Likelihood (see *Materials & Methods: Assigning USVs* and *Discussion* for details).

HyVL performed significantly better than either method alone, allowing a total of 91.1% of USVs to be assigned at a spatial accuracy of 4.8 mm (MAE). This constitutes a substantial, 2.9-fold improvement in accuracy over the previous state of the art, the SLIM algorithm.^73^ On the full set of USVs where both microphone arrays were recording (N = 7982), HyVL outperformed both USM4/SLIM and Cam64/Beamforming significantly, both in residual error (SLIM: 14.8 mm; Cam64: 5.33 mm; HyVL: 5.07 mm; p<10^-10^ for all comparisons, Wilcoxon rank sum test) and percentage of reliably assigned USVs (SLIM: 74.4%; Cam64: 80%; HyVL: 91.1%). Cam64/Beamforming performed even more precisely on its reliably assignable subset (4.56 mm), which was, however, smaller than the HyVL set. This difference emphasizes the complementarity of the two methods and thus the synergy through their combination. There was no significant difference between tracking on dyadic and triadic recordings (HyVL: 5.0 mm vs. 5.1 mm, p=0.71, Wilcoxon rank sum test) with correspondingly similar selection percentages (92 vs. 90%, respectively).

The accuracies above are an average over localization performance at any distance. In particular during close interaction, USVs will often be reflected or obstructed, complicating localization. While this constitutes the realistic challenge during mouse social interactions, we also investigated the ‘ideal’, unobstructed performance of HyVL by comparing the performance on USVs emitted when all animals were ‘far’ (>100 mm) apart, i.e. more than ~20 times the average accuracy of HyVL, as well as for a single male mouse on the platform. For the *far* USVs the reliably assignable fraction increased to 97.9%, and the accuracy significantly improved to 3.8 mm (*Fig*. 3C gray, p=8.6×10^-7^, Wilcoxon rank sum test). For the *single animal* USVs, the accuracy was even better at 3.4 mm with 98.4% reliably assigned (*Fig*. 3C, blue).

Next, we inspected separate localization along the X and Y axis to check for anisotropies of localization (*Fig*. 3D/E, histograms normalized to maximum). The position of the closest animal aligned precisely with the estimated position in both dimensions, indicated by the high density along the diagonal *(Pearson r* > 0.99 for both dimensions) and the MAE’s along the X and Y direction separately (X = 3.1 mm, Y = 2.8 mm). These one-dimensional accuracies might be of relevance for interactions where movement is restricted.

Lastly, we visualized the localization density relative to the mouse that the vocalization was assigned to (*Fig*. 3F). Combining both dimensions and appropriately rotating them, the estimated position of the USVs is shown relative to the mouth. The density is narrowly centered on the snout of the mouse (circle radius = MAE: green: SLIM method; orange: HyVL; light orange: HyVL assigned USVs, gray: Far assigned USVs).

In summary, the HyVL system provides a substantial improvement in the localization precision. In comparison to other methods, its precision also allows a larger fraction of vocalizations to be reliably assigned and retained for later analysis, which enables a near complete analysis of vocal communication between mice or other vocal animals.

### Sex Distribution of Vocalizations During Social Interaction

Courtship interactions between mice lead to high rates of vocal production, but are challenging due to the relative proximity, including facial contact. Previous studies using a single microphone have often assumed that only the male mouse vocalized,^84–87^ while more recent research has concluded that female mice vocalize as well.^76,88^ Female vocalizations were typically less frequent, but constituted a substantial fraction of the vocalizations (11-18%). Below, we demonstrate that the accuracy of the localization system can be an important factor for conclusions about the contribution of different sexes to the vocal interaction.

Overall, female vocalizations constitute the minority of vocalizations overall. Naive estimation without MPI selection using SLIM estimates ~14%, while HyVL tallies it at just 7% (Fig. 4A). Applying MPI selection, SLIM estimates only 5.5%, while HyVL arrives at significantly less, just 4.4% (p=0.002, paired Wilcoxon signed rank test, Fig. 4A/B), while reliably classifying 91.1% of all vocalizations.

**Figure 4:**
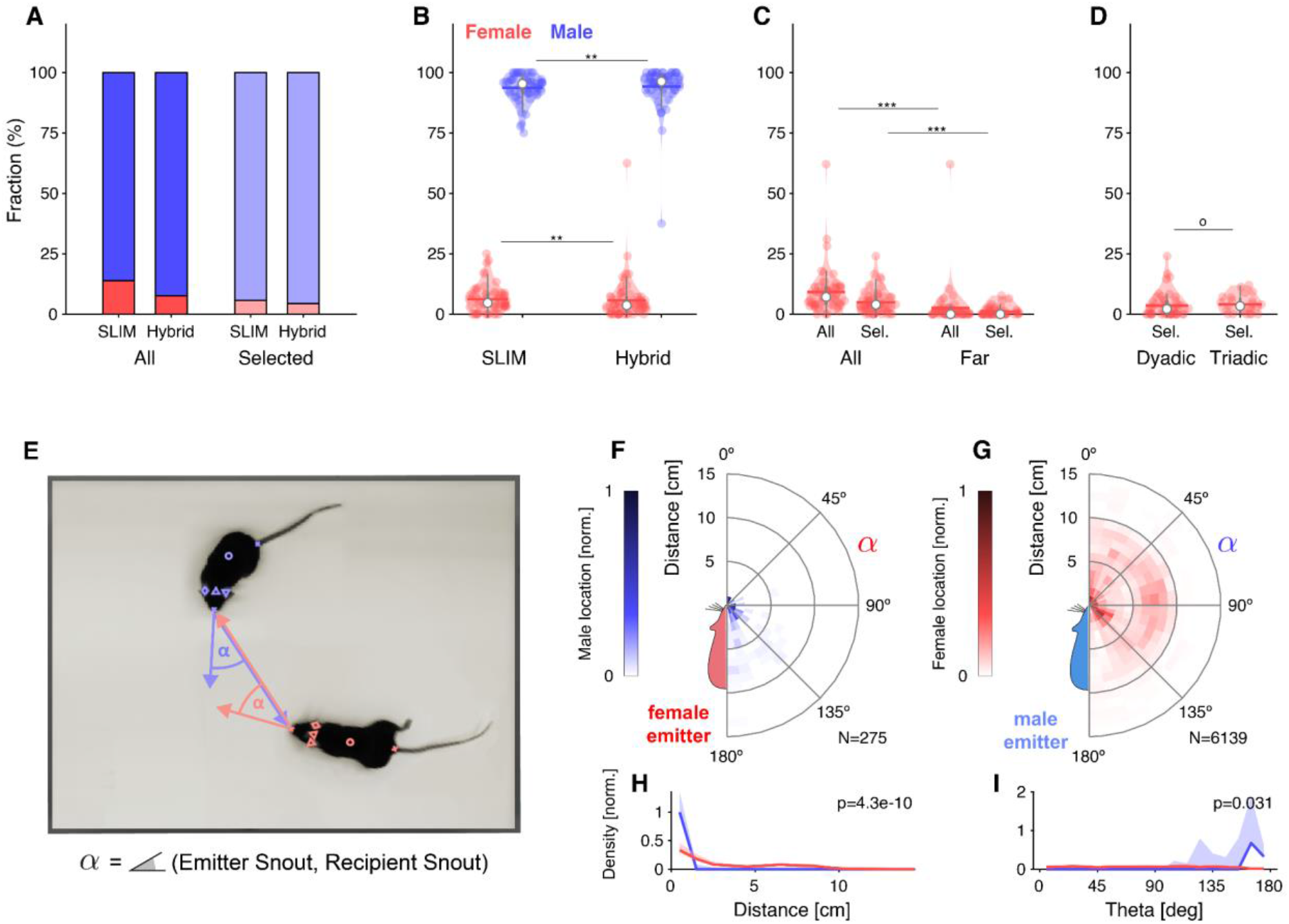
Analysis of Sex-dependent Vocalizations can depend on Localization Accuracy. **A** Female vocalizations constitute a small fraction of the total set of vocalizations. The female fraction further reduces with increased precision and when selecting vocalizations based on the MPI. **B** Using the hybrid method instead of SLIM significantly reduces the fraction of female vocalizations, suggesting that less accurate algorithms overestimate the female fraction (only results for MPI-selected USVs shown). **C** The fraction of female vocalizations further reduces if only USVs are considered that are emitted while all animal snouts were >50 mm apart from each other. This indicates a preference of female mice to vocalize in close snout-snout contact, however, this entails that female vocalizations are more prone to confusion with male vocalizations due to their relative spatial occurrence. **D** There was no difference in the female fraction of USVs between dyadic and triadic pairings (2 male and 2 female conditions combined here). **E** High-accuracy localization of USVs allows one to analyze the relative spatial vocalization preferences of the mice, i.e. their occurrence density in relation to the relative position of other mice to the emitter. We quantified this by collecting the position of the non-vocalizing mice at the times of vocalization, in relation to the vocalizing mouse. ***α*** corresponds to the angle between the emitter’s snout and the snout of other mice. **F** Female mice appear to emit vocalizations in very close snout-snout contact, with a small fraction of vocalizations also occurring when the male mouse around the hind-paws/anogenital region. **G** Male mice emit vocalizations both in snout-snout-contact, but also at greater distances, which dominantly correspond to a close approach of the male’s snout to the female anogenital region. This was verified separately with a corresponding analysis, where the recipient’s tail-onset was used instead (not shown). **H** Radial distance density of receiver animals, marginalized over directions, shows a significant difference, with females vocalizing mostly when males (blue) are in close proximity of the snout, while males vocalize when the female mouse’s snout is very close (corresponding to snout-snout contact), but also when the female’s snout is about 1 body length away (snout-anogenital interaction). Plot show medians and percentile-based confidence bounds. **I** Direction density of receiver animals, marginalized over distances, shows borderline significance with female mice vocalizing mostly when the male snout is located behind their own snout.

Using HyVL instead of SLIM significantly reduces the fraction of female vocalizations, suggesting that less accurate algorithms overestimate the female fraction (only results for MPI-selected USVs shown, Fig. 4B). Considering only vocalizations that are emitted when the snouts are >50 mm apart, further significantly reduces the fraction to female USVs to 1.1%after MPI selection (p=5.2×10^-8^, Wilcoxon Rank Sum test). Comparing the percentage of female vocalizations between dyadic and triadic trials, no significant differences were found (p=0.22, Wilcoxon Rank Sum test, Fig. 4D).

Beyond the absolute distance between the mouths of the mice, high-accuracy localization of USVs allows one to position the bodies of the animals relative to one another at the times of vocalization by combining acoustic data with multiple concurrently tracked visual markers. This provides an occurrence density of other mice relative to the emitter (Fig. 4E).

Female mice appear to emit vocalizations in very close snout-snout contact, with a small fraction of vocalizations occurring when the male snout is around the hind-paws/anogenital region (Fig. 4F). Male mice emit vocalizations both in snout-snout-contact, but also at greater distances, which dominantly correspond to a close approach of the male’s snout to the female anogenital region (Fig. 4G). This was verified separately with a corresponding analysis, where the recipient’s tail-onset was used instead (not shown).

In summary, the combination of high-precision localization and selection using the MPI indicate that female vocalizations may be even less frequent than previously thought. When they vocalize, the mice appear to almost exclusively be in close snout-snout contact. As this is incidentally also the condition which has the highest chance of mis-assignments, even the remaining female vocalizations need to be treated with caution.

### Vocalization Rate Analysis

In dyadic trials, one female and one male mouse interacted, whereas in triadic trials either two males and one female or two females and one male mouse interacted. In the analysis of triadic interactions, we separate competitive and alternative contexts depending on whether a mouse had to compete with another same sex mouse or could interact with two opposite sex mice, respectively. For triadic trials we further separate the same-sex mice into dominant and subordinate, based on who vocalized more.

Overall vocalization rates did not differ significantly between dyadic and triadic conditions for male and female mice (see *Fig*. 5A). However, in competitive interactions between males, one male mouse significantly and strongly dominated the ‘conversation’, with on average 9-fold more vocalizations than the other male mouse (T_D_ vs T_s_, both comparisons: p<0.0001, Wilcoxon Sum of Ranks test; retains p<0.05 after Bonferroni correction for multiple testing, *Fig*. 5A,B). While the present division into dominant and subordinate mouse based on a higher vocalization rate within a recording will always lead to a significant difference, the quantitative difference between them is the striking aspect in this comparison. Overall male vocalization rates were similar in competitive and alternative triadic trials. Female vocalization rates were similar across all compared conditions.

**Figure 5:**
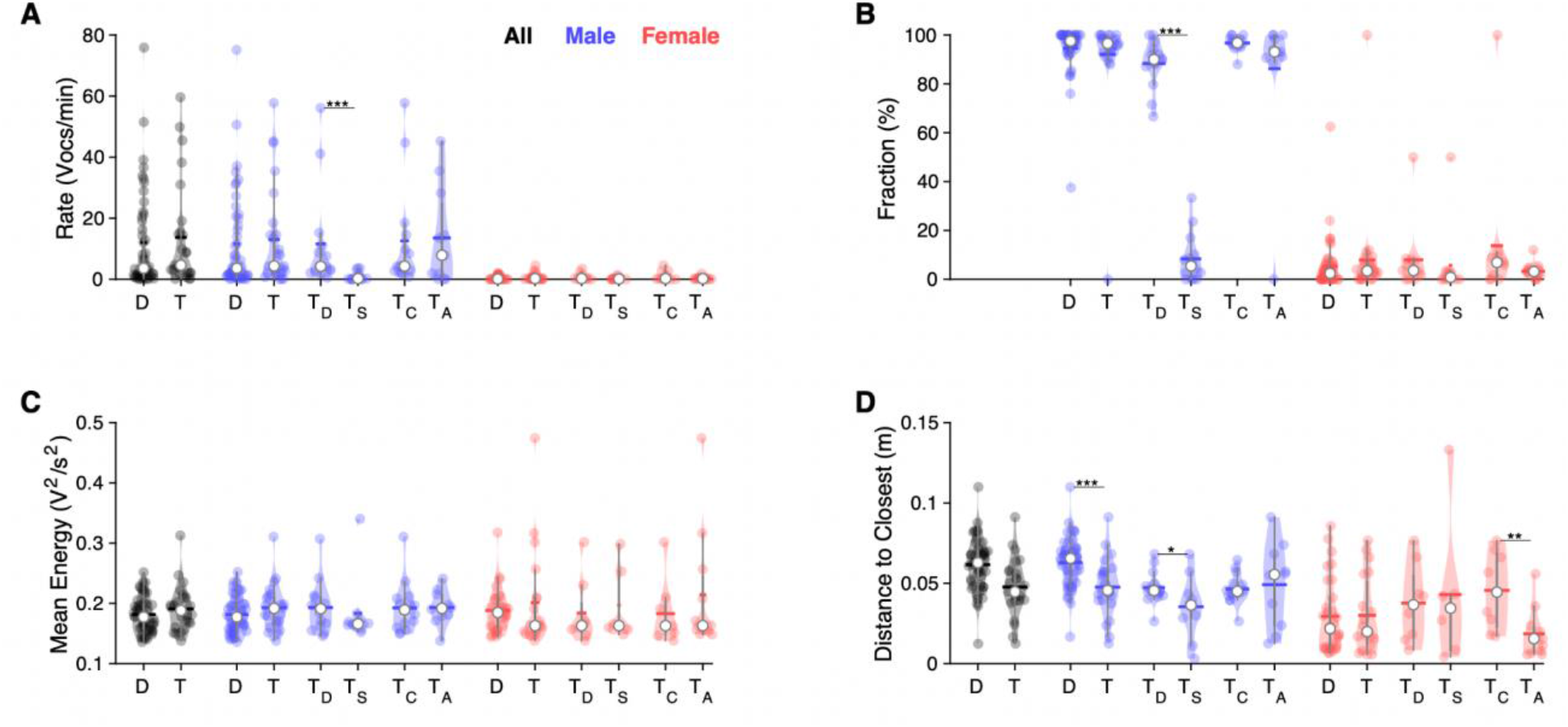
In triadic interaction, one male vocalizes dominantly and males vocalize even closer to females. **A** Overall, vocalization rates were comparable between dyadic (D) and triadic (T) conditions. Male mice (blue) vocalized at higher rates than female mice (red). However, this was restricted to the dominant male mouse (T_D_: dominant = emitted more USVs within same-sex) in triadic, competitive (2m/1f) conditions (see text for all p-values.). Male vocalization rates were similar in competitive (T_C_: with same-sex competitors;) and alternative (T_A_: no same-sex competitor, i.e. for male vocs: 2f/1m) pairings. Female vocalization rates remained low and similar across all conditions. T_S_: submissive mouse = emitted fewer USVs within same sex during competitive trial; white dot: median; horizontal bar: mean. **B** While the fraction of USVs emitted by males was overall comparable between D and T pairings, the dominant male (T_D_) emitted a substantially larger fraction than their submissive counterpart (T_S_), roughly a factor of 9. In competitive pairings, male mice tended to emit an overall larger fraction of all USVs than in alternative pairings (T_C_ vs. T_A_), but this is unsurprising as both males vocalize. In female mice, the overall fraction of USVs in D and T pairings was also similar (see details in Results for potential caveats of the dominant/subordinate classification). **C** In triadic pairings, dominant male mice tended to vocalize more intensely than in dyadic pairings, however, this difference was not significant at the current sample size. No significant differences were found for female mice. **D** Male mice emitted USVs in closer proximity to the closest female mouse in triadic compared to dyadic interactions. Female mice generally emitted USVs at closer distances (see also Fig. 4F/H), in particular for alternative vs. competitive pairings.

The mean vocalization energy of dominant males in triadic pairings tends to be higher than those of submissive males in triadic pairings, however, this result did not reach significance in the present dataset (see *Fig*. 5C). No effects of vocalization energy were found in females.

The distance to the closest animal of opposite sex was found to be even closer during triadic trials (see *Fig*. 5D), driven purely by male vocalizers (p=0.0003, Wilcoxon Sum of Ranks test): the distance to the closest animal does not change between conditions for vocalizing females (p=0.975, Wilcoxon Sum of Ranks test). Interestingly, the distance to the closest animal was larger for females at the time of vocalization when they had a same-sex competitor on the interaction platform with them than when they were the only female (Tc vs. Ta, p=0.0068, Wilcoxon Sum of Ranks test).

In summary, in competitive triadic interactions, one of the male mice took a strongly dominant role and approached the female even more closely when vocalizing than in dyadic pairings. The latter suggests that more vocalizations occurred during snout-snout interactions than in dyadic interactions, where the bulk was in snout-to-anogenital interactions. Correspondingly, female mice also vocalized more snout-snout in alternative interactions from their perspective, pointing to overall more vocalizations during snout-snout interactions when two male mice competed for a female mouse.

## Discussion

We have developed and evaluated a novel, hybrid sound localization system (HyVL) for ultrasonic vocalizations (USVs) emitted by mice and other rodents. USVs are innately used by rodents to communicate social and affective information and are increasingly being used in neuroscience as a behavioral measure in neurodevelopmental and neurolinguistic research. In the context of dyadic and triadic social interactions between mice, we demonstrate that HyVL achieves a groundbreaking increase in localization accuracy down to ~3.4-4.8 mm, enabling the reliable assignment of >90% of all USVs to their emitter. Further, we demonstrate that this can be combined with automatic tracking, enabling a near-complete and automated analysis of vocal interaction between rodents. The showcased analyses demonstrate the advantages obtained through more precise localization, further discussed below. HyVL is based on an array of high-quality microphones in combination with a commercially available, affordable acoustic camera. With our freely available code, this system can be readily reproduced by other researchers and has the potential to revolutionize the study of natural interactions of mice.

### Comparison with previous approaches for localizing vocalizations

Localization accuracy was first systematically reported by Neunuebel et al.^76^ using a 4-microphone setup and a maximum likelihood approach,^89^ who attained an MAE of ~38 mm that conferred an assignment rate of 14.6-18.1% (their Table 1, *assigned* relative to *detected* or *localized)*. Originating from the same research group, Warren et al.^75^ employed both a 4 and 8 microphone setup in a follow-up study, achieving an MAE of ~30 mm for 4 microphones (~52% assignment rate) and of ~20 mm with 8 microphones (~62% assignment rate), both using a jackknife approach to increase robustness of localization. Stahl et al.^73^ introduced the SLIM algorithm, reaching an MAE of ~11-14 mm (~80-85% assignment rate depending on the dataset) using 4 microphones. Presently, we advance the state-of-art in multiple ways: we use 68 microphones, combining a 64 channel ‘acoustic camera’ with 4 high quality ultrasonic microphones. While the acoustic camera has relatively basic micro-electromechanical systems (MEMS) microphones, it is inexpensive, features a high degree of integration and correspondingly easy operation. Combining the complementary strengths of the two arrays is the key advantage of the present approach over previous approaches, as it allows for a quantum leap in accuracy (3.4-4.8 mm, 91% assignment rate), while keeping the complexity of the system manageable. A comparable alternative might be a 16-channel array from high-quality microphones, which would, however, be substantially more expensive (~€40,000) as well as cumbersome to build and refine.

A future generation of MEMS microphones might make the use of the high-quality microphones unnecessary and thus further simplify the system setup, allowing for inexpensive, small-form factor deployment (see below).

### Expected impact for future research

Mice and rats are social animals,^90,91^ and isolated housing^92^ or testing^93^ can affect subsequent research outcomes. Social isolation also has direct effects on the number and characteristics of USVs, at least in males.^94,95^ Sangiamo et al.^88^ demonstrated that distinct USV patterns can be linked to specific social actions and the latter that locomotion and USVs influence each other in a context-dependent way. Using HyVL, such analyses could be extended to more close-range behaviors, when a substantial fraction of the vocalizations are emitted (see *Fig*. 1D). The development of more unrestricted behavioral paradigms, made viable by increased localization precision, will thus also likely prove valuable to the fields of human language impairment and animal behavior. As an added benefit, better USV localization will also likely increase lab animal well-being via (i) more social contact in specific cases where they spend much time with their conspecifics in the testing environment, or when the home environment is the testing environment (e.g., PhenoTyper; Noldus Information Technologies), and (ii) a reduced need for (non-)invasive markers.

Here, we conducted a limited set of showcase analyses on the spatial characteristics of vocalization behavior. As expected, the system was accurate enough to assign vocalizations during many snout-snout interactions as well as other, slightly more distant interactions, e.g. snout contact with the ano-genital region of the dyadic partner. We found the male mice to vocalize most while making snout contact with the abdomen and ano-genital region of the female wild-type. Females vocalized predominantly during snout-snout contact, with the male approaching from behind.

This highlights an example of how localization accuracy can shape our understanding of roles in social interaction between mice: A recent, pivotal study^76^ demonstrated that female mice vocalize during courtship interactions. Research from our group^73^ concluded further that mice primarily vocalize in snout-snout interactions, incidentally the condition that makes assignment the most difficult. While the present results maintain that female mice vocalize, the fraction appears to be lower than previously thought. We, however, emphasize that this conclusion still requires further study under different social contexts, e.g. interaction of more mice as in some of the previous studies.^34,88^

The compact form factor of the HyVL microphone arrays, in particular the Cam64, enables studies of social interaction in home cages. There, rodents are less stressed and likely to exhibit more natural behavior, in particular if the home cage includes enrichments. The relatively low hardware costs for HyVL allows deployment of multiple systems to cover larger and more natural environments.

### Current limitations and future improvements of the presented system

The millimeter accuracy by HyVL enables the assignment of USVs even during close interaction, certainly including all snout-anogenital interactions, and many snout-snout interactions. However, certain snout-snout interactions are still too close to reliably assign co-occurring USVs. While the MPI criterion maintains reliability even then, subsequent analysis will be partially biased due to the exclusion of these USVs during the closest interactions. While a further improvement of accuracy may be possible, close inspection of the sound density maps available via beamforming from the Cam64 recordings suggests that the mouse’s snout acts as a distributed source: the sound density is rather evenly distributed on it, without a clear internal peak. During free interaction, we noticed that the sound density was co-elongated with the head-direction of the mouse and could thus be used as an additional feature to identify the vocalizer. However, this proved unreliable during close interaction, likely due to absorption and reflection of sounds based on the mice’s bodies. More advanced modeling of the local acoustics or deep learning might be able to resolve these issues, but would require ground truth recordings, which could be obtained in interactions where one mouse is known to be silent, e.g. by cutting the vocal cords.

The present strategy for combining the estimates from Cam64/Beamforming and USM4/SLIM was chosen as it optimized the reliably assigned percentage of USVs, while minimizing the residual distance. We also tested alternative approaches, e.g. using direct beamforming on the combined data from Cam64 and USM4 (unreliable estimates, due to mismatch of number of microphones, not further pursued), maximum likelihood combination of estimates (MAE=7.1 mm),^96^ and making the selection solely depend on the MPI (MAE=5.2 mm). While each of these approaches have certain, theoretically attractive features, the results were worse in each case, likely due to particular idiosyncrasies of the MPI computation, the different microphone characteristics, and the estimation of single-estimate uncertainty.

Lastly, for the purpose of online feedback during experiments and to reduce data warehousing, it would be advantageous to perform the localization of USVs in real-time. This would be enabled by streaming the data to a GPU, performing localization immediately and keeping only a single channel, beamformed estimate of each USV. Ideally, the same device could run visual tracking simultaneously, which would remove all temporal limitations on the recordings in terms of data size and enable continuous audiovisual tracking.

### Conclusion

HyVL delivers breakthrough accuracy and assignment rates, likely approaching the physical limits of assignment. The low system costs (<€10k) in relation to its performance make HyVL an excellent choice for labs studying rodent social interaction. Many recent questions regarding the sequencing of vocalizations during social interactions become addressable with HyVL without intrusive interventions. Its use can both refine the precision and reliability of the analysis, while reducing the number of animals required to complete the research due to a larger fraction of assigned USVs per animal.

## Materials & Methods

All experimental procedures were approved by the animal welfare body of the Radboud University under the protocol DEC-2017-0041-002 and conducted according to the Guidelines of the National Institutes of Health.

### Animals

In our experiment, 4 female C57Bl/6J-WT, 6 male C57Bl/6J-WT and 8 male C57Bl/6J-Foxp2.flox/flox;L7-cre mice (bred locally at the animal facility) were studied. For subsequent analyses, WT and KO mice were combined (see beginning of Results for reasoning). The mice were 8 weeks old at the start of the experiments. After 1 week of acclimation in the animal facility, the experiments were started. Mice of the same sex were housed socially (2 to 5 mice per cage) on a 12-hour light/dark cycle with ad libitum access to food and water in individually ventilated, conventional EU type II mouse cages at 20°C with paper strip bedding and a plastic shelter for basic enrichment. Upon completion of the experiments, the animals were anesthetized using isoflurane and sacrificed using CO_2_.

The current experiment was performed as an add-on to an existing set of experiments, whose focus included a region-specific knockout of *Foxp2* in the cerebellar Purkinje cells of the male mice, denoted as C57Bl/6J-Foxp2.flox/flox;L7-cre. Neither previous work nor our own work has detected any differences in USV production between WT and KO animals^97^, so - given the mostly methodological focus of the present work - we considered it acceptable to pool them in the current analysis, reducing the number of animals needed, thus treating all males as WT C57Bl/6J, the genotype of the female mice.

### Recording Setup

The behavioral setup consisted of an elevated interaction platform in the middle of an anechoic booth together with 4 circumjacent ultrasonic microphones as well as an overhanging 64-channel microphone array and high-speed video camera (see *Fig*. 1A).

The booth had internal dimensions of 70 x 130 x 120 cm (L x W x H). The walls and floor were covered with acoustic foam on the inside (thickness: 5 cm, black surface Basotect Plan50, BASF). The acoustic foam shields against external noises above ~1 kHz with a sound absorption coefficient > 0.95 (N.B., defined as the ratio between absorbed and incident sound intensity), which corresponds to >26 dB of shielding apart from the shielding provided by the booth itself. In addition, the foam strongly attenuates internal reflections of high-frequency sounds like USVs. Illumination was provided via 3 dimmable LED strips mounted to the ceiling, providing light from multiple angles to minimize shadows.

The support structure for the interaction platform and all recording devices was a common frame constructed from slotted aluminum (30×30 mm) mounted to the floor of the anechoic booth, guaranteeing precise relative positioning throughout the entire experiment. The interaction platform itself was a 40×30 cm rectangle of laminated, white acoustic foam (thickness 5 cm; Basotect Plan50) chosen to maximize the visual contrast with the mice and simplify the cleaning of excreta. The interaction platform had no walls to avoid acoustic reflections and was located centrally in the booth. Its surface was elevated 25 cm above the floor (i.e. 20 cm above the foam on the booth floor), which was generally sufficient in preventing animals from leaving the platform. If a mouse left the platform, data was excluded from further analysis (<5% of frames).

Sounds inside the booth were recorded with 2 sets of microphones: (i) 4 high-quality microphones (USM4) and (ii) a 64-channel microphone array (Cam64), both recording at a sampling rate of 250 kHz at 16 bits. (i) The 4 high-quality microphones (CM16/CMPA48AAF-5V, AviSoft, Berlin) were placed in a rectangle that contained the platform (see *Fig*. 1A) at a height exceeding the platform by 12.1 cm to minimize the amount of sound blocked by the mice during interaction. The position of a microphone was defined as the center of the recording membrane. Considering the directional receptivity of the microphones (~25 dB attenuation at 45°), the microphones were placed a short distance away from the corners of the platform to maximize sound capture (5 cm in the long direction and 6 cm in the short direction of the platform). The rotation of each microphone was chosen to be such that it aimed at the platform center. The microphones produce a flat (±5 dB) frequency response within 7-150 kHz that was low-pass filtered at 120 kHz to prevent aliasing (using the analog, 16th order filter, which is part of the microphone amplifier). Recorded data was digitized using a data acquisition card (PCIe-6351, National Instruments). (ii) In addition, a 64-channel microphone array (Cam64 custom ultrasonic version, Sorama B.V.) was mounted above the platform with a relative height of 46.5 cm measured to the bottom of the Cam64 and a relative lateral shift of 6.52 cm to the right of the platform midpoint. The Cam64 utilizes 64 MEMS microphones (Knowles, Digital Zero-Height SiSonic, SPH0641LU4H-1) for acoustic data collection that are positioned in a Fermat’s spiral over a circle with a ~16 cm diameter. Raw microphone data was streamed to an m.2 SSD for later analysis. Synchronization between the samples acquired by the Cam64 and the ultrasonic microphones was performed by presenting 2 brief acoustic clicks (realized by stepping a digital output from 0 to 5V) close to one of the microphones on the Cam64 at the start and end of each trial using a headphone driver (IE 800, Sennheiser). The recorded pulses were automatically retrieved and used to temporally align the recording sources.

A high-speed camera (PointGrey Flea3 FL3-U3-13Y3M-C, Monochrome, USB3.0) was mounted above the platform with a relative height of 46.5 cm measured to the bottom of the front end of the lens (6 mm, Thorlabs, part number: MVL6WA) and a relative lateral shift of 4.48 cm to the left of the platform midpoint. Video was recorded with a field of view of 52.2 x 41.7 cm at ~50 fps and digitized at 640 x 512 pixels (producing an effective resolution of ~0.815 mm/pixel). The shutter time was set to 10 ms to guarantee good exposure while keeping the illumination rather dim. The frame triggers from the camera were recorded on an analog channel in the PCIe-6531 card for subsequent temporal alignment with the acoustic data.

### Experimental Procedures

The experiment had 3 conditions: dyadic (with 2 mice; 57 trials), triadic (with 3 mice; 28 trials), as well as monadic (single male mouse, ground truth data; 8 trials). Two dyadic trials were excluded from further analysis due to repeated but required experimenter interference during the recordings, leaving 55 dyadic trials. Each trial consisted of 8 minutes of free interaction between at least 1 female and at least 1 male mouse on the platform. Females were always placed on the platform first, and males were added shortly thereafter. In the monadic case, fresh female urine was placed on the platform to prompt the male mouse to vocalize. The high-speed camera and 4 high-quality microphones started recording after all mice had been placed on the platform and continued for 8 minutes. Data points where one mouse had left the platform or the hand of the experimenter was visible 10 seconds before or after (e.g., to pick up a mouse) were discarded (<5% of frames). Due to the rate of data generation of the Cam64 recordings (32 MB/s), their duration and timing was optimized manually. The experimenter had access to the live spectrogram from the USM4 microphones, and upon the start of USVs, triggered a new Cam64 recording (of fixed 2 min duration). If additional USVs occurred after that point, the experimenter could trigger additional recordings.

### Data Analysis

The analysis of the raw data involved multiple stages (see *Fig*. 2): From the audio data, the presence and origin of USVs was estimated automatically. From the video data, mice were carefully tracked by hand at the temporal midpoint of each USV as a ground truth comparison for their acoustically localized origin. To estimate what proportion of our precision would be lost when using a faster and more scalable visual tracking method, we also tracked the mice automatically during dyadic trials. The estimated locations of the mice and USVs were then used to attribute the USVs to their emitter. All these steps are described in detail below.

#### Audio Preprocessing

Prior to further analysis, acoustic recordings were filtered at different frequencies. USM4 data was band-pass filtered between 30-110 kHz before further analysis using an inverse impulse response filter or order 20 in Matlab (function: designfilt, type: bandpassiir). Cam64 data was band-pass filtered with a frequency range adapted to the frequency content of each USV. Specifically, first the frequency range of the USV was estimated as the 10th to 90th percentile of the set of most intense frequencies at each time point. Next, this range was broadened by 5 kHz at both ends, and then limited at the top end to 95 kHz. If this range exceeded 50 kHz, the lower end was set to 45 kHz. This ensured that beamforming was conducted over the relevant frequencies for each USV and avoided the high-frequency regions where the Cam64 microphones are dominated by noise (see *Fig*. 1C).

#### Video Preprocessing

The high-speed camera lens failed to produce perfect rectilinear mapping and was placed off-center with respect to the interaction platform, thereby producing a nonlinear radial-tangential visual distortion. We corrected for the radial distortion with:

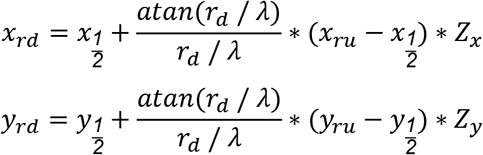

where [*x_rd_, y_rd_*] represent the radially distorted image coordinates, 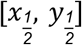 the coordinates of the image center, *r_d_* the Euclidean distance to the radial distortion center, *λ* the distortion strength, [*x_ru_, y_ru_*] the radially undistorted coordinates, and *Z_x_, Z_y_* axis-specific zoom factors. The tangential distortion, on the other hand, we corrected with:

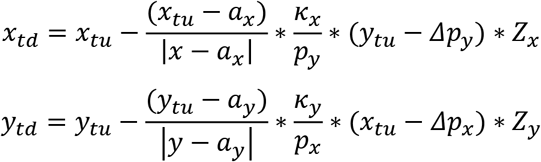

where [*x_td_, y_td_*] represent the tangentially distorted image coordinates, [*x_tu_, y_tu_*] the tangentially undistorted coordinates, [*a_x_, a_y_*] the coordinates of the tangential distortion center, [*x, y*] the size of the image, [*κ_x_, κ_y_*] the tangential distortion strengths, [*p_x_, p_y_*] the size of the interaction platform in the undistorted image, and [*Δp_x_, Δp_y_*] the offset of the platform with respect to the top-left corner of the undistorted image.

#### Detection of Ultrasonic Vocalizations

USVs were detected automatically using a set of custom algorithms (see `VocCollector.m`) described elsewhere.^72^ Detection was only performed on the USM4 data, as their sensitivity and frequency range was generally better than for the Cam64 (see *Fig*. 1C). A vocalization only had to be detected on 1 of the 4 high-quality microphones to be included into the set. In total, we collected 13406 USVs, out of which 8424 occurred when the Cam64 recordings were active.

#### Automatic Visual Animal Tracking

To assess whether we could reliably assign USVs to their emitter in a fast and scalable way, we automatically tracked multiple body parts of interacting mice — most importantly the snout and head center — for all dyadic trials (see *Fig 2*). The tracking task can be separated into 2 steps: (i) identifying all candidate locations for different body parts in the video and (ii) linking these body part locations over time without switching identities between the animals. We performed the first step by analyzing the video data offline in the XY-plane with *DeepLabCut*, a toolbox that uses a convolutional neural network to visually identify animal features (DLC; see Mathis et al., 2018). To train the network, a training set was used (~1400 frames) containing manually placed markers for both mice (i.e., snout, tail base, head center, left/right ear, body center). The training set was constructed in 2 iterations, with the problematic aspects of training using the first iteration (i.e., ~800 randomly drawn frames) prompting frame choice in the second iteration (i.e., ~600 manually selected frames). Subsequently, DLC was trained on the basis of this data (DLC v.2.2b8, running on a GTX 1070 GPU with NVIDIA driver version 390.77 on Ubuntu 18.04.1 LTS; see *Supplementary Material* for a comprehensive overview of all DLC parameters). The resulting convolutional neural network was then used to predict the marker locations in all frames and all trials and provided quite high spatial precision, although not all visible markers were represented by a location estimate in every frame. In addition, while more recent versions of DLC (v.2.2+) should also be able to track identities consistently over time, we did not manage to achieve high reliability in that respect using DLC. Taken together, every trial would have required hundreds of manual corrections to achieve reliable tracking over the entire period without identity switches.

To solve these 2 problems, we developed a dedicated solution in MATLAB (see `C_trackMiceDLC.m`) that used spatial location estimates generated by DLC but disregarded their unreliable identity. The general strategy was as follows: first, we requested in DLC not just the estimate with the highest probability rating for each marker but rather an estimate cloud of the 50 most likely candidate locations for each marker and each mouse (see *Suppl. Fig*. 1A). Within each marker class, all estimates were treated as identical at this point. Optimal marker locations were then generated by within-frame k-means clustering of the estimate clouds (N.B., with *k* being varied in a data-driven manner), or if clustering was suboptimal, by probability-weighted averaging of heuristically separated estimate clouds (see *Suppl. Fig*. 1B and `LF_BPExtract` in `C_trackMiceDLC.m`). After that, all markers underwent a temporal and spatial analysis in tandem, both aimed at constructing pieces of unattributed tracks. In the temporal analysis, marker positions that obviously belonged together were first assembled into small spatiotemporal threads with the same, unknown identity (i.e., belonging to the same mouse). The speed and acceleration that these small threads represented were subsequently assessed over time to yield longer threads of marker positions with the same identity (see *Suppl. Fig*. 1C and `LF_tempAttr` in `C_trackMiceDLC.m`). In the spatial analysis, all marker positions were analyzed on a frame-by-frame basis, grouping markers with the same identity based on a logical combination of anatomically permitted inter-marker distances (see *Suppl. Fig*. 1D and `LF_spatAttr` in `C_trackMiceDLC.m`). Next, recognizing the overlap between the unattributed threads and the attributed sets of marker positions, the spatial and temporal information was logically combined in its entirety. This process was mathematically equivalent to a discrete convolution of the sets of integers representing marker identities with the discrete response function describing the thread to which they belong. For example, if a snout and head center clearly belong together in a specific frame, the threads that run through them also belong together (see *Suppl. Fig*. 1E and `LF_attrConv` in `C_trackMiceDLC.m`). Afterwards, the thread ends still loose were connected based on quadratic spatial trajectory estimates for each marker individually. Lastly, all marker trajectories were smoothed with piece-wise shape-preserving cubic interpolation. Reliable tracking was ensured for all recordings by visual inspection and, if required, manual corrections (~10 per trial on average, a major reduction).

#### Manual Visual Animal Tracking

We manually tracked the spatial locations of all mice during all USVs from the video data to assess the precision of the automatic visual and acoustic tracking. During manual tracking, the observer was presented with a combined display of the vocalization spectrogram and the concurrent video image at the temporal midpoint of each USV *(MultiViewer*, custom-written, MATLAB-based visualization tool). The display included a zoom function for optimal accuracy, as tracking was click-based. Users could also freely scroll in time to ensure consistent animal identities. Only the snout and head center (i.e., midpoint between the ears) needed to be annotated because these points define a vector representing the head location and direction, which was all that was required in subsequent behavioral analyses.

#### Localization of Ultrasonic Vocalizations

USVs were spatially localized using a hybrid approach that drew on the complementary strengths of the 2 microphone arrays (see *Fig*. 2). For example, the Cam64 array provided excellent localization for USVs with energy below ~90 kHz, due to the increasing noise floor of the MEMS (microelectro-mechanical systems) microphones with sound frequency. Conversely, the 4 high-quality ultrasonic microphones (USM4) have a rather flat noise level as a function of frequency. On the other hand, USM4 will occasionally have glitches in one of the microphones, which can be compensated for in Cam64-based estimates through the number of microphones. As a consequence, the errors of the two methods show an L-shape (see *Fig*. 3A), which highlights the synergy of a hybrid approach.

Acoustic localization using the Cam64 recordings was performed on the basis of delay-and-sum beamforming^82^. In beamforming, signals from all microphones are combined to estimate a spatial density that correlates with the probability of a given location being the origin of the sound. Specifically, we computed beamforming estimates for a surface situated 1 cm above and co-centered with the interaction platform, extending to 5 cm beyond all edges of the platform (i.e., 50 x 40 cm in total) at a final resolution of 1 mm in both dimensions. We refer to this density of sound origin as *D_so_*(*x, y*) where *x* and *y* denote spatial coordinates. To prevent noises unrelated to a specific USV from contaminating the location estimate, we limited beamforming to a particular frequency range estimated from the simultaneous data of the USM4 array that enveloped the USV. Spatial density was defined as

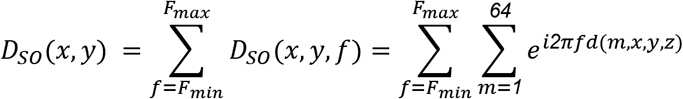

where *d*(*m*, *x*, *y*, *z*) denotes the difference in arrival time at each microphone *m* for sounds emitted from a location with coordinates (*x,y,z*), where *z* is omitted in *D_so_*(*x,y*) as it is a fixed distance to the plane of the microphone array. Beamforming was performed in the computational cloud backend provided by the Cam64 manufacturer, the so-called Sorama Portal^1^.

The final beamforming estimate was calculated sequentially in 2 steps: first, a coarse estimate with 1 cm resolution was generated over the entire beamforming surface. Second, a fine-grained estimate with 1 mm resolution was generated over a 30 x 30 mm window centered on the peak location of the coarse estimate (see *Fig*. 2 for an example). This two-step approach was chosen to optimize performance, as an estimate with 1 mm resolution over the entire beamforming surface would be computationally expensive while failing to produce a better result.

For USVs of sufficient quality (i.e., containing frequency content below ~90 kHz while being sufficiently intense and long), both the coarse and fine estimates of *D_so_*(*x,y*) contained a peak whose height was typically very large compared to the surrounding values at distances greater than a few cm’s. The peak location of the fine-grained estimate was used as the final estimate of the USV’s origin. To assess the quality of this location estimate, we computed a signal-to-noise ratio (SNR) per USV as follows:

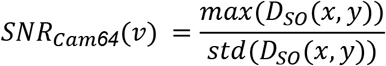

where *D_so_*(*x,y*) is assumed to be calculated for the USV *ν*. The inverse, *1/SNR_Cam64_* was used as a proxy for the uncertainty of localization for a given USV.

Localization from the USM4 recordings was performed using the SLIM method^73^. Briefly, SLIM analytically estimates submanifolds (in 2D: surfaces) of a sound’s spatial origin for each pair of microphones and combines these into a single estimate by intersecting the manifolds (in 2D: lines). The intersection has an associated uncertainty which scales with the uncertainty of the localization estimate for a given USVs, specifically the uncertainty was defined as the standard deviation of all locations that were >90% times the maximum of the intersection density of all origin curves.

Lastly, for each USV where both Cam64 and SLIM location estimates *Ẋ_Cam64_* and *Ẋ_SLIM_* were available, a single estimate *Ẋ_HyVL_* was computed based on the two estimates, spatial uncertainties and their spatial relation to the mice at the current time (see below).

#### USV Assignment

The final, hybrid location estimate and assignment to a mouse was performed while taking into account the probability of making a false assignment as proposed before^76^, through the calculation of the mouse probability index *MPI*. While the *MPI* was previously only used to exclude uncertain assignments (e.g. if two mice are nearly equidistant to the estimated sound location), we also adapted it here to select and combine the location estimates. The *MPI_k_* for each mouse *k* was computed as

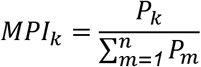

Here, *P_k_* is the probability that the USV in question originated from mouse *k* computed as 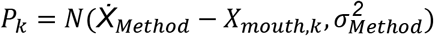, where *Ẋ_Method_* is an estimate of the acoustic origin, *Ẋ_mouth, k_* the position of the mouth of mouse *k*, and 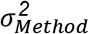 the uncertainty of the estimate, with *Ẋ_Method_* and 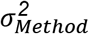 specific to the Method used. *X_mouth,k_* was assumed to lie on a line connecting the snout and head-center. For manually tracked recordings, the optimal location on this line was close to the snout (~2% towards the head, where % is relative to the snout-to-head-center tracked distance), while in the automatic tracking it was ahead of the snout tracking point (~15% away from the head). 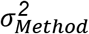 was computed for each USV as the method’s intrinsic per-USV uncertainty estimate. As these uncertainty estimates only correlate with the absolute uncertainty (i.e. in mm’s), we scaled them such that their average across all USVs matched the residual error of each method in the Far-condition (all animals >100 mm apart, see *Fig*. 3C and Oliveira-Stahl et al.^73^). In this way, the *MPI_k_* for individual USVs took into account the uncertainty of each method: if the uncertainty of one method was higher, probabilities across mice would become more similar and the *MPI_k_* would reduce.

For a given USV, we computed the *MPI_k_* for all mice for both methods. The mouse with the largest *MPI_k_* per method - which coincides with the mouse at the smallest distance to the estimate - was denoted as *MPI_Cam64_* and *MPI_SLIM_*, respectively. If only one of the two exceeded 0.95, this method’s estimate was selected. If both exceeded 0.95, then the estimate with the smaller distance to the mouse with the highest *MPI_k_* was chosen. This combination ensured that only reliable assignments were performed, while minimizing the residual error. Similar to Neunuebel et al. 2015, we also excluded estimates that were too far away from any mouse (50 mm). This distance threshold mainly serves to compensate for a deficiency of the *MPP*. if all mice are far from the estimate, all *P_k_* are extremely small, however, the *MPI_k_* will often exceed 0.95. The distance threshold corresponds to setting the individual *P_k_* = 0 in the *MPI_k_*, thus excluding candidate mice which are highly unlikely to be the source of the USV. USVs that had no *MPI_k_* > 0.95 for either method were excluded from further analysis. The fraction of included USVs is referred to as *selected* in the plots. Maximizing this fraction is essential to perform a complete analysis of vocal communication.

We compared the above-described combination strategy to a large number of alternative strategies, including maximum likelihood combination of estimators (Ernst & Banks 2002), or selecting directly based on the largest *MPI_k_* or largest *P_k_*. While all these approaches led to broadly similar results, the described approach achieved the most robust and reliable results (see Discussion for additional details).

#### Audiovisual Alignment

For both microphone sets, precise measurements of their location in relation to the camera’s location were used to position acoustic estimates in the coordinate system of the images provided by the camera. In the final analysis, we noticed for each microphone set small, systematic (0.5-2 mm) shifts in both X and Y. We interpreted these as very small measurement errors in the relative positions of the camera or microphone arrays, and corrected these post-hoc in the setup definition, followed by rerunning all subsequent analysis steps. This reduced all systematic shifts to near 0.

#### Spatial Vocalization Analysis

To gain insight into the spatial positioning of the interacting mice, we represented the relative animal positions in a polar reference frame centered on the snout of the emitter. In this format, the radial distance corresponded to the snout-snout distance and the radial angle described the relative angle between the gaze direction of the emitter and the snout position of the recipient (i.e., with the line from the head center to the snout of the emitter pointing towards 0°; see also *Fig*. 5A).

The position density of the recipient mouse was collected in cumulative fashion, with the polar coordinate system translated appropriately for each USV based on its temporal midpoint. We assumed that the mice had no preference for relative vocalizations to either side of their snout, so all relative spatial positions were agglomerated in the right hemispace for further analysis. All data points were then binned using a polar, raw-count histogram with bins of 10° and 1 cm.

#### Statistical Analysis

To avoid distributional assumptions, all statistical tests were nonparametric, i.e., Wilcoxon rank sum test for two-group comparisons and Kruskal-Wallis for single factor analysis of variance. Correlations were computed as Spearman’s rank-based correlation coefficients. Error bars represent standard errors of the mean (SEM) unless stated otherwise. All statistical analyses were performed in MATLAB v.2018b (The Mathworks, Natick) using functions from the Statistics Toolbox.

## Supporting information

Suppementary Methods

## Acknowledgements

We would like to thank Lucas Noldus for suggesting the use of the Sorama Cam64 and Maurice Camp and Toros Senan for technical support relating to the operation and data handling of the Cam64 and the Sorama Portal. We would like to thank Amber van der Stam, Dionne Lenferink, Soha Farboud for assisting with the animal handling and experimental control. BE acknowledges funding from a DCN Internal Grant, funded by Noldus IT b.v. as well as an NWO VIDI grant (016.VIDI.189.052) and a Technology Hotel Grant (ZonMW, 40-43500-98-4141).

## Footnotes

1 www.sorama.eu/sorama-portal

## Notes

**Competing Financial Interests:** The authors declare that they do not have any competing financial interests associated with the present work.

### Competing Interest Statement

The authors have declared no competing interest.

