## Supplementary material for "Rodent ultrasonic vocal interaction resolved with millimeter precision using hybrid beamforming": Suppementary Methods

for

M. L. Sterling[1], B. Englitz[1]

[1]Computational Neuroscience Lab, Donders Institute for Brain, Cognition and Behaviour, Radboud University, Nijmegen, The Netherlands

**Supplementary Methods:**

*Automatic Animal Tracking Precision*

To assess how a faster and more scalable tracking method would impact our localization precision relative to manual tracking, we also performed visual mouse tracking automatically during dyadic trials. With this approach, tracking is not temporally restricted to the midpoint of USV production, but can be performed for every frame of the entire recording. This data can be used to establish spatial densities of interaction against which e.g. the spatial density of vocalizations can be compared.[54]

Mice were tracked offline using a combination of *DeepLabCut (DLC)*[65] and extensive post-processing to maintain animal identity over the entire recording. While the tracking results from DLC were generally quite accurate, we refrained from using them directly because of inaccuracies and identity switches that occurred on many hundreds of occasions in every recording. Instead we adopted a strategy where DLC generated an overcomplete set of candidate locations followed by custom synthesis and tracing of these alternatives in space and time (see *Materials & Methods: Automatic Visual Animal Tracking*; and *Suppl. Fig*. 1). In short, improved marker locations were generated from marker estimate clouds produced by DLC. Next, these marker positions were assembled into short spatiotemporal threads with the same, unknown identity based on a combination of spatial and temporal analysis. Finally, the thread ends were connected based on quadratic spatial trajectory estimates for each marker, yielding the complete track for both mice. This strategy resulted in reliable, high-quality tracking for all recordings, with a greatly reduced number of manual corrections needed overall (~10 per trial on average). All resulting tracks were visually verified (representative example: *supplementary video 1*).

We compared the accuracy of localization on the basis of manual tracking with that of automatic tracking (N = 5046 USVs, see *Suppl. Fig.* 2). Directly comparing the snout positions between the methods shows a median difference of 3.76 mm. The resulting error for localizing USVs was still superior to other systems, however, significantly increased by ~0.9 mm (MAE = 5.71 mm) relative to manual tracking. Both manual and automatic tracking appear to have particular patterns of residual errors, indicated by the fact that the error between the tracking methods is much larger than their difference in USV localization error. The percentage of reliably assignable USVs interestingly increased to 93.6% (HyVL), compared to 92% with manual tracking for the dyadic recordings only. We optimized the mouth location on the snout-to-head-center line, finding an optimal distance of 15% of the snout to head center distance *to the front* of the animal. This indicated that the automatic tracking tended to place the snout tracking point a bit further into the snout than manual tracking, which might also explain the increase in assignment, due to a slight - but erroneous - increase in the separation between the snouts. While these results suggest that manual tracking is still advantageous, it highlights that completely automatic analysis of dyadic and possibly n-adic social interaction experiments is feasible at slightly reduced accuracy.

Below, we showcase two analyses that are enabled by the high accuracy of HyVL, where the focus is on their technical feasibility.

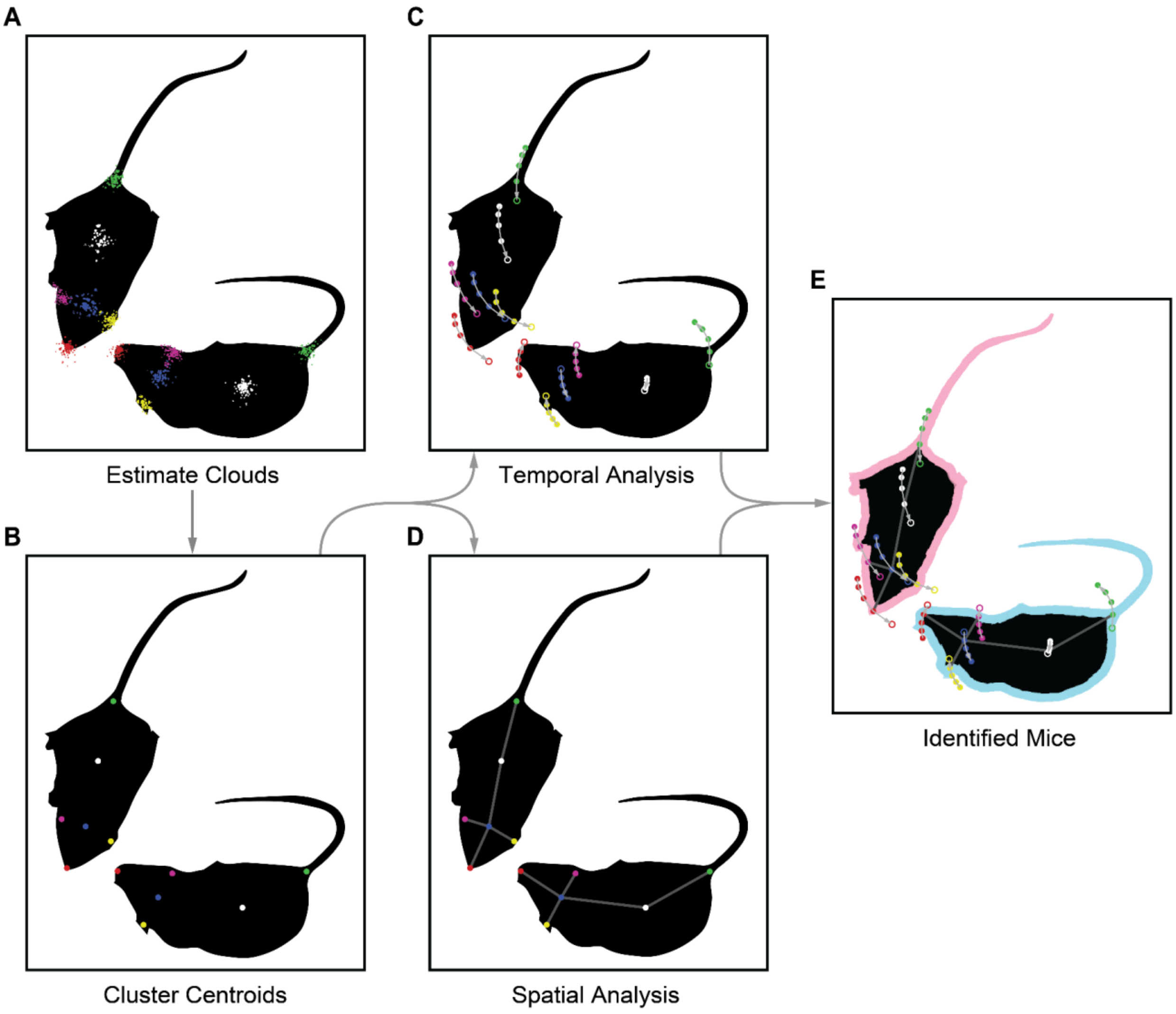


**Supplementary Figure 1:** Schematic depiction of the progression of marker localization and identity attribution.

**A** First, unattributed estimate clouds (i.e., not assigned to an animal) for each of the six markers are collected using DeepLabCut (DLC). Within each marker class, all estimates are treated as identical at this point (colors indicate marker class).

**B** Optimal marker locations are then generated by within-frame, k-means clustering of the estimate clouds, or if that results in an incorrect number of clusters, by probability-weighted averaging of estimate clouds heuristically separated by animal.

**C, D** Next, all markers undergo a temporal and spatial analysis in tandem, both with the goal of constructing pieces of unattributed tracks.

**C** In the temporal analysis, small spatiotemporal threads of marker locations are assembled and assessed in terms of speed and acceleration to extend them as much as possible across neighboring frames.

**D** In the spatial analysis, all marker positions are analyzed spatially on a frame-by-frame basis, grouping markers with the same identity based on a logical combination of anatomically permitted inter-marker distances.

**E** Finally, complete, attributed tracks are constructed by combining both analyses (see *Methods: Automatic Visual Animal Tracking*).

For actual data, the analysis is less ideal than in the figure: not all markers necessarily have an estimate cloud representing them all the time, nor are the estimates always accurate. The finished tracks were visually checked and corrected if necessary (~10 corrections per trial on average, a major reduction).

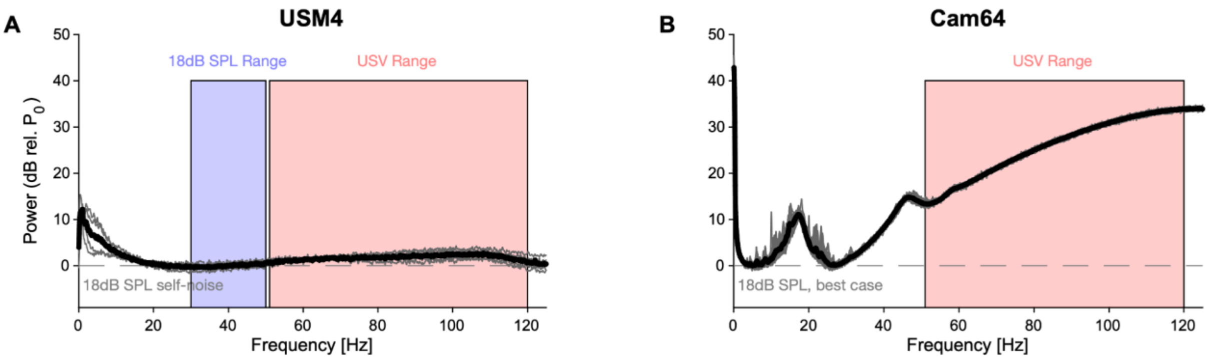


**Supplementary Figure 2:** Comparison of noise spectra of the two microphone arrays. Spectra are computed from the output voltage of the microphones during the same silent stretch of a recording (duration: 1s, M82,R41, T = [150,151]s), verified to not contain any discernible vocalizations, footsteps or other discrete sounds.

**A** The high-quality ultrasonic microphones (gray: 4x individual, black: average) had a rather flat baseline-noise level across frequency, with the only exception of an ~10 dB deviation for low frequencies, far below the relevant frequency range contained in USVs (red overlay). For reference, the spectrum was shifted to the average of the power in the 30-50 kHz range (blue overlay), which is listed in the manual as being the "input-referred self-noise level", corresponding to 18 dB SPL[1].

**B** The Cam64 MEMS microphones (gray: 64x individual, black: average, Akustica AKU242[2]) have a highly frequency-dependent baseline-noise across frequency. The peaks at frequencies < 50 kHz are less relevant for USV detection, while the substantial rise of the noise level for higher frequencies poses an issue for detecting very high frequencies, as their contribution to the recorded signal gets progressively buried in the baseline noise (see Fig. 1C for the effect on the detectability of high-frequency USVs). Since no information was available on the input-referred self-noise level in the technical documentation, we shifted the curve to its minimum. In reality it should be shifted higher to be quantitatively compared with the Avisoft microphone, as the latter's large membrane is expected to outperform the AKU242 at all frequencies. For clarity, the above spectra are not equivalent to the sensitivity of the microphone at different frequencies, however, the baseline noise limits the sensitivity at these frequencies. While in principle a frequency dependent increase in sensitivity could overcome the baseline noise, this does not seem to be the case (see Fig. 1C, top).

[1] http://www.avisoft.com/ultrasound-microphones/cm16-cmpa/
[2] https://nl.mouser.com/datasheet/2/720/PB24-1.0%20-%20AKU242%20Product%20Brief-770082.pdf

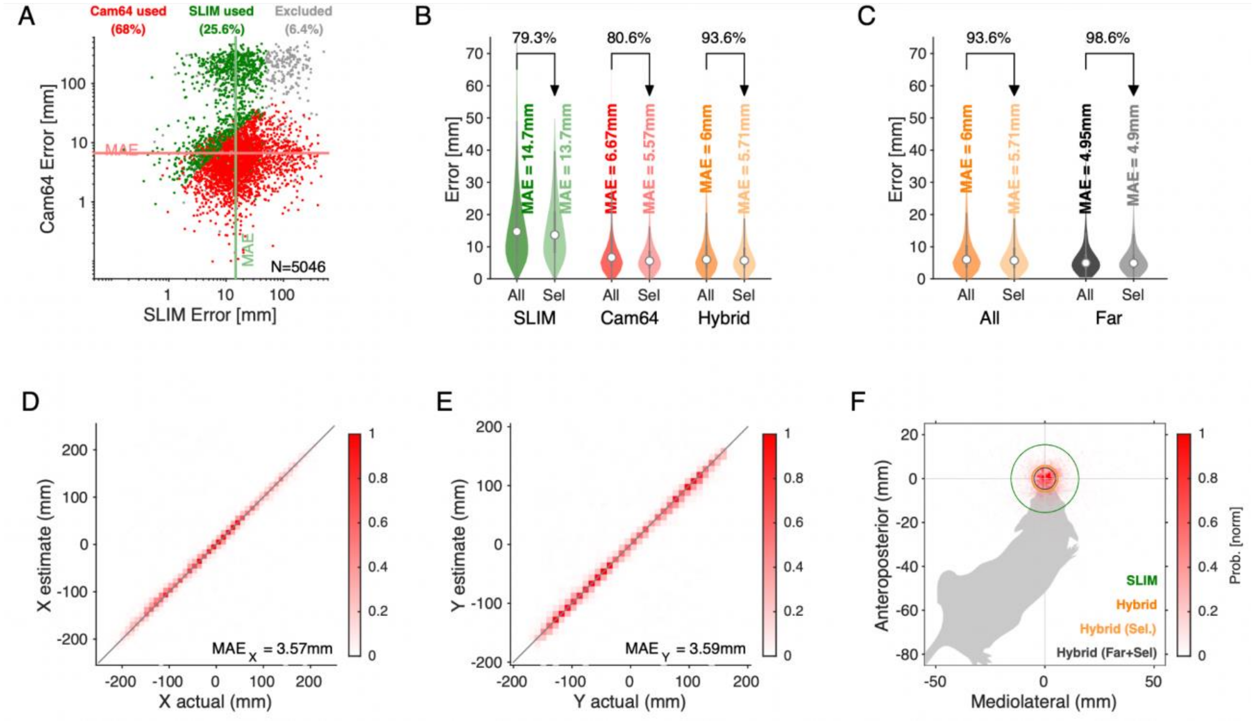


**Supplementary Figure 3:** Localization results on the basis of automatic tracking from dyadic and triadic recordings. All plotting matched to the manual results for dyadic and triadic tracking in *Fig.* 3. In panel A, the scaling of the axes is linear instead of logarithmic as in Figure 3.

**Supplementary Video 1:** Example of HyVL tracking and sound localization

Marker color represents animal sex (light blue: male; light red: female). Marker shape represents body part (circle: body center; cross: snout or tail; downward triangle: left ear; upward triangle: head center; diamond: right ear). Cam64 USV localizations (yellow) are overlaid on the beamforming densities (red) which are often very narrow and therefore hard to see underneath the localization marker (yellow dot). SLIM USV localizations are shown as well (orange '+'), typically further away from the snout in comparison to Cam64-based localization markers.
